# Science Family skills: An Alexa Assistant Tailored for Laboratory Routine

**DOI:** 10.1101/484147

**Authors:** Tiago Lubiana-Alves, André A.N.A. Gonçalves, Helder I Nakaya

## Abstract

Voice User Interfaces such as Amazon Alexa and Google Home are already widely available and used for personal purposes. These services could be used to improve experimental biology laboratory routine, facilitate troubleshooting and increase efficiency. Till date, no applications that are tailored to enhance laboratory routine have been made available. Here, we present a set of free-to-use, open source tools adapted to Alexa for application in the laboratory environment, with prospects of enhancing productivity and reducing work-related stress. All skills, 3D printer model and source codes are freely available in the Alexa app store and in GitHub.

## INTRODUCTION

Alexa is Amazon’s Voice User Interface (VUI) that utilizes cloud-based artificial intelligence to deliver, among other things, information on weather, traffic, and Wikipedia, as well as read out aloud the appointments from your calendar or just play your favorite songs. Although the introduction of technology to laboratory routine is on the rise^1^, VUI devices are present in millions of homes across the globe^2^, and their applications to experimental science have been explored^3^, these virtual assistants are not routinely used inside a laboratory.

We have created a set of custom-developed applications (i.e., “skills”) that allow Alexa reply to your voice command and read out loud a protocol, the latest news, and articles from a given subject or tell you the precise location of a specific laboratory reagent (Figure 1). More importantly, through the use of voice commands, scientists can keep doing research without taking off their gloves.

**Figure 1.**
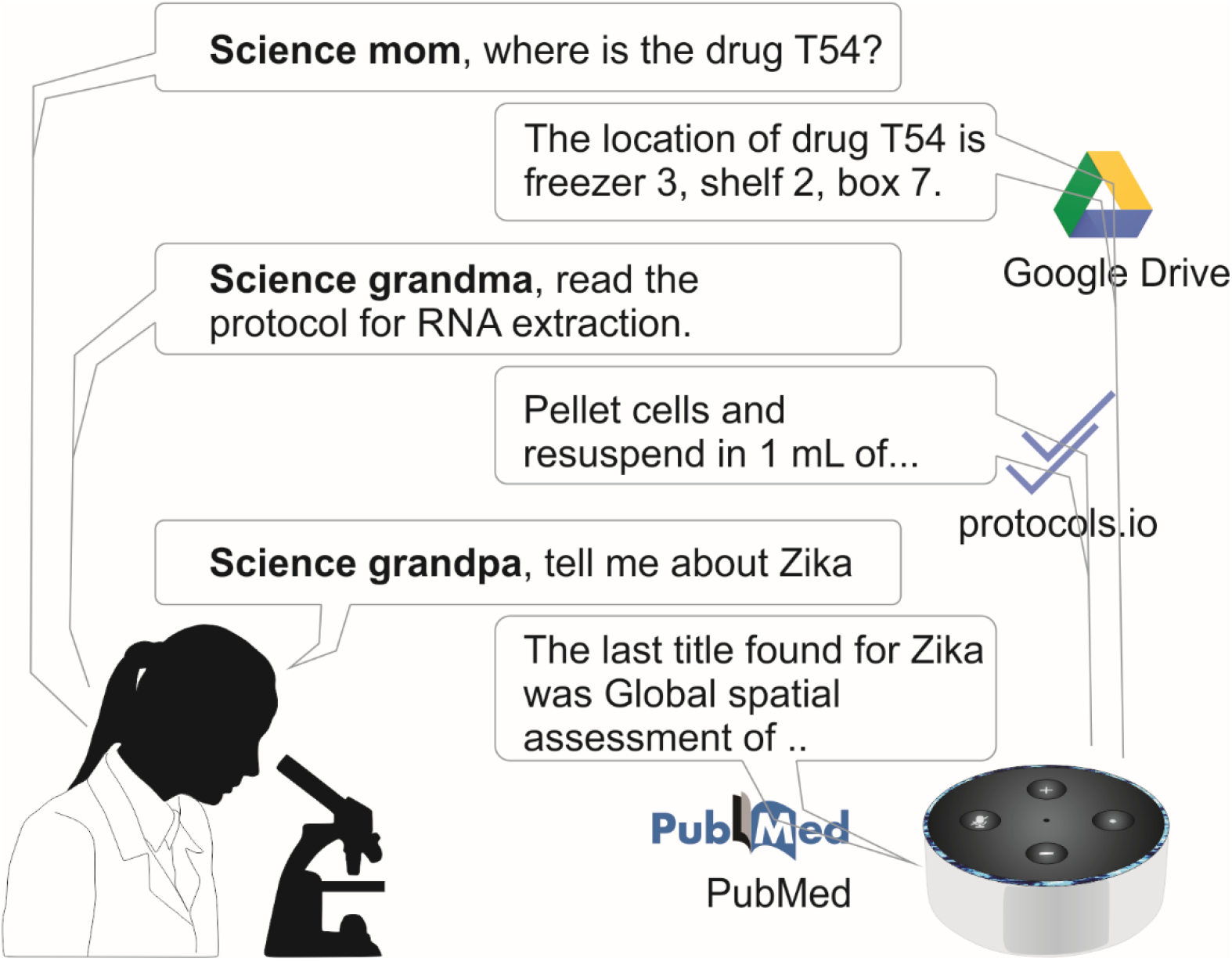
Scheme highlighting the process of using the Science Family Alexa skills in the laboratory.

To facilitate development and community support, we have made its functionalities distinct. The “Science Family” suite contains four applications at the moment, and each skill should be download separately from Amazon Alexa app store.

### Science Mom skill

“Science Mom”^4^ is designed to localize items (reagents and equipment) in the laboratory. After uploading the laboratory inventory into a Google sheet, users may ask questions such as “where can I find glove boxes?” or “where is the restriction enzyme box?”, and Science Mom will inform them of their location. Since the single sheet can be shared with all members of a laboratory, everyone may edit the inventory, updating Alexa about the location of every object in the lab.

### Science Grandma skill

“Science Grandma”^5^ reads out loud any protocol the same way grandmas read recipes. This skill is integrated with a user-friendly and collaborative database of online protocols^6^, *https://www.protocols.io/*. This database allows the access to protocol details of a plethora of ready-made laboratory procedures as well as custom protocols. The Protocols.io IDs can be inserted in a Google Sheet, just as in Science Mom, and mapped to a user-defined name. Then, it is possible to make requests such as “start reading the RNA Extraction protocol” and use commands such as “go to next step” or “go to step five” to navigate through the protocol.

### Science Grandpa skill

“Science Grandpa”^7^ provides an on-demand query of biomedical literature by sourcing the PubMed database. If you ask questions such as “what are the news about Malaria?” or “which are the most relevant articles on Chaga’s Disease?”, Science Grandpa will access PubMed and retrieve relevant (according to the database algorithm) or recent publications on these topics. You can skip the articles by saying “next” while hearing the titles. If a given article title interests you, say “send,” and Alexa will send the article’s abstract directly to your email.

### Alexa Lab Holder

We also developed a 3D printer model for the Alexa echo dot device and made the model template available in a public repository^8^. The printed object can hold the Alexa echo dot device vertically on a laboratory shelf through the use of a printed screw (Figure 2). This allows researchers to place Alexa anywhere within the lab.

**Figure 2.**
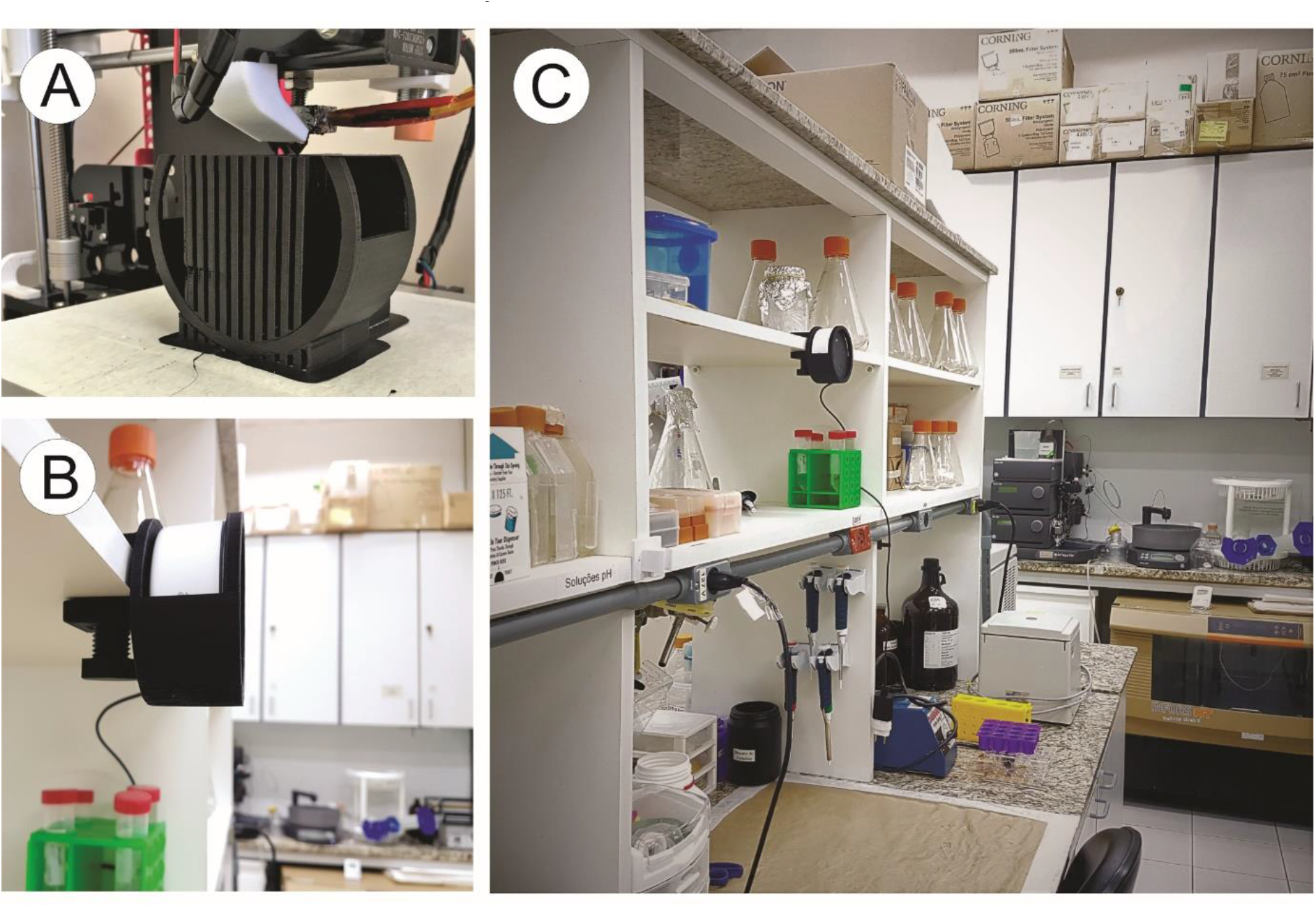
Alexa Lab holder 3D model. (a) We used the Autodesk Fusion 360 software (educational license) to do the 3D modeling of Alexa holder. The Alexa holder was made using the 3D Anet A8 printer, with layer resolution of 0.2mm and 10% of infill. The raw material for printing was Polylactic acid (PLA) and the total printing time was 7 hours and 40 min. (b) Side view showing Alexa Lab holder docked on a laboratory shelf. (c) Distant view of Alexa Lab Holder inside a research laboratory.

## CONCLUSIONS

We present the first publicly-available VUI targeted to help experimental researchers. The user-friendliness and low-cost of the hardware needed for it makes it readily accessible to virtually any laboratory in the world. All source codes are freely available in GitHub^9^.

It is important to note that our source code is modular, allowing for a parallelized development while, at the same time, blocking the side effects of modifications in one branch on the others. More importantly, it is open-source, making it possible for the community to improve and customize the project. Scientists whose native languages are not English can also readily translate the code and make the skills available in any Alexa-supported languages (such as Japanese, German, Spanish, French, and Italian).

The employment of this suite of applications will lead to the creation of a more productive work environment, facilitating the adaptation of new students and providing fast troubleshooting for laboratory routine. We envision that additional skills may be included into the Science Family to allow a straightforward execution of even more complex tasks, such as ordering new reagents, controlling lab equipment, and recording experimental data.

## ACKNOWLEDGMENTS

We wish to thank Maysa Braga Barros Silva, Maryana Stephany Ferreira Branquinho, Maria Carmen Sale e Silvana Sandri for helpful reviews and essential feedback.

## AUTHOR CONTRIBUTIONS

T.L. wrote the software; T.L and H.I.N. designed the software and contributed to the final manuscript; A.G. designed the Alexa Lab Holder 3D model; H.I.N. supervised the execution of the project.

